# Aberrant chromatin looping by NUP98-HOXA9 is constrained by CTCF and facilitated by cohesin

**DOI:** 10.64898/2026.06.22.733801

**Authors:** Tooba Rashid, Zachary A Drum, Ivana Y. Quiroga-Barber, Doris Cruz Alonso, Gelila Petros, Eric Davis, Benjamin Kornegay, Chelsea Yang, Chenxi Xu, Sylvie Parkus, Gang Greg Wang, Wesley R Legant, Jill M Dowen, Douglas H Phanstiel

## Abstract

The acute myeloid leukemia (AML) fusion protein NUP98-HOXA9 (NHA9) drives leukemogenesis by promoting aberrant chromatin loop formation through phase separation, yet the mechanisms underlying these interactions remain unclear. To address this, we dissect the interplay between NHA9 and individual loop extrusion factors using in situ Hi-C, CUT&RUN, RNA-seq, and Auxin-inducible degradation of CTCF or RAD21. CTCF was found to be dispensable for NHA9 loop formation, although CTCF binding constrained a subset of loops that emerged only upon CTCF depletion. In contrast, cohesin played a distance-dependent role where short-range NHA9 loops formed independently of RAD21, while long-range loops were strongly cohesin-dependent. Despite this requirement, RAD21 showed minimal enrichment at NHA9 loop anchors, indicating that NHA9 does not function as a canonical cohesin barrier. Instead, these findings support a non-canonical model in which cohesin transiently facilitates interactions between distal NHA9-bound loci, which are subsequently stabilized through NHA9 phase separation. Together, this work reveals a distinct mechanism of oncogenic chromatin looping in which NUP98-HOXA9 cooperates with canonical loop extrusion machinery to reprogram genome architecture in AML.

---

Chromatin looping is a central mechanism by which distal regulatory elements communicate with gene promoters to control transcription^1–3^. In mammalian cells, the majority of chromatin loops are thought to arise through loop extrusion, a process mediated by the cohesin complex and constrained by the architectural protein CTCF. Cohesin extrudes chromatin until it encounters convergently oriented CTCF sites, establishing a structural framework for gene regulation^4–9^. While this model explains the vast majority of chromatin loops found in human cells, recent studies have uncovered several non-canonical mechanisms of chromatin loop formation^10–12^.

We previously showed that the oncogenic fusion protein NUP98–HOXA9 (NHA9), which is associated with a high-risk subtype of acute myeloid leukemia, drives the formation of aberrant chromatin loops through mechanisms that differ from canonical loop extrusion^12^. NHA9 arises from a chromosomal translocation that fuses the intrinsically disordered FG-repeat domain of NUP98 with the DNA-binding homeodomain of HOXA9, enabling it to bind specific genomic loci and promote their coalescence into phase-separated nuclear condensates^12,13^. These interactions preferentially connect H3K27ac-marked regulatory elements with promoters of leukemia-associated genes, including HOX, PBX3, and MEIS1, and are associated with increased gene expression and cellular proliferation. Notably, the anchors of these loops are largely not bound by CTCF, distinguishing them from canonical loop extrusion-mediated interactions. Conversely, cells expressing a phase-separation-deficient NHA9 mutant fail to form these loops, do not upregulate target gene expression, and do not exhibit increased proliferation^12^.

Despite these observations, the relationship between NHA9-mediated looping and canonical genome organization remains unclear. In particular, it is unknown whether architectural proteins such as CTCF and cohesin play any functional role in the formation, maintenance, or spatial organization of these non-canonical loops. Resolving this question is important not only for understanding how distinct mechanisms of genome organization are integrated but could also inform the future development of therapeutic interventions that target these aberrant chromatin structures.

Here, we systematically investigate the role of loop extrusion machinery in NHA9-mediated chromatin looping using *in situ* Hi-C, CUT&RUN, RNA sequencing, and inducible degradation of CTCF and cohesin. We find that NHA9 establishes chromatin loops independently of CTCF anchoring but within a genome architecture that is constrained by CTCF-defined domain boundaries. Surprisingly, NHA9 loops are largely dependent on cohesin in a distance-dependent manner, with longer loops exhibiting a stronger dependence on RAD21. RAD21 does not accumulate at NHA9 loop anchors, suggesting that NHA9 does not function as a cohesin barrier in the way that CTCF does. Instead, we propose that cohesin-driven loop extrusion brings NHA9 binding sites together initially, but phase separation of NHA9 then stabilizes and maintains these interactions.

## Results

### NUP98-HOXA9 forms chromatin loops in HCT116 cells

To study the role of CTCF and cohesin in NHA9-driven chromatin looping, we employed the Auxin-inducible degron (AID) system, which enables rapid and selective degradation of endogenously tagged proteins upon Auxin treatment. We employed two HCT116-derived cell lines, each containing AID tags for either CTCF or RAD21 (a core component of the cohesin complex). Each cell line was engineered to stably express mCherry-tagged NHA9 using the PiggyBac transposon system. We performed *in-situ* Hi-C on NHA9-expressing cells and their corresponding parental cell lines, including treatments with DMSO and Auxin (1.74 billion average reads across 8 conditions). Comparison of NHA9-expressing and parental cells revealed widespread changes in chromatin looping, including the formation of 109 gained loops and 33 lost loops (DESeq2, padj < .1, absolute log2FC > 0; **Fig. 1A–B**). These results indicate that NHA9 expression induces *de novo* chromatin looping in HCT116 cells, consistent with our previous observations in HEK293T cells^12^.

**Fig. 1.**
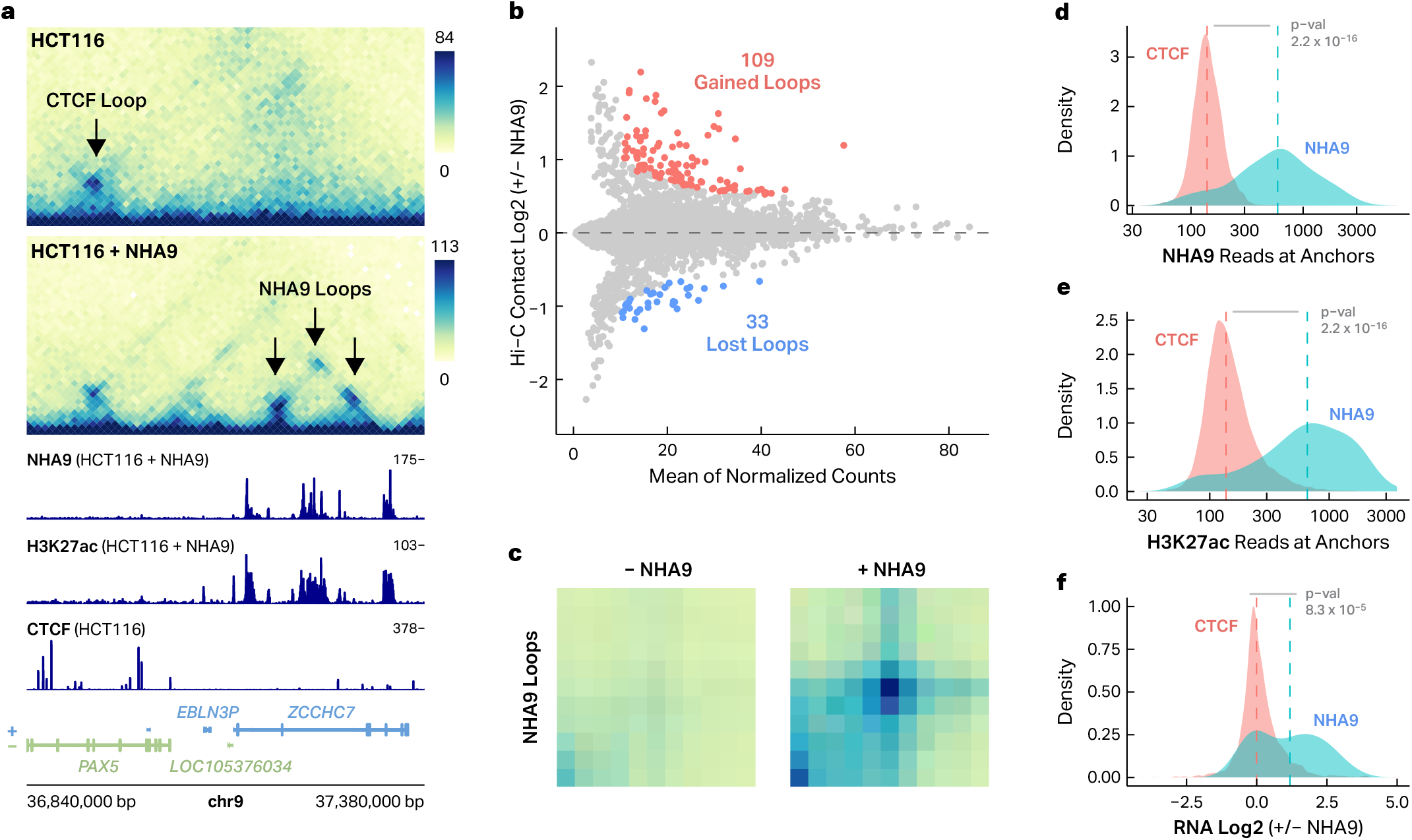
NUP98-HOXA9 forms chromatin loops in HCT116 cells. **a** Representative region in HCT116 cells that do not and do express NHA9, with labeled arrows indicating CTCF and NHA9 loops. Signal tracks for NHA9, H3K27ac, and CTCF. The cell line and genotypes are indicated in parentheses. **b** MA plot of all chromatin loops comparing HCT116 cell lines that do not and do express NHA9. Differential loops are colored in red and blue (DEseq2, padj < 0.1, absolute log2 fold change > 0). **c** Aggregate Peak Analysis (APA) plots of annotated NHA9 loops in HCT116 cells that do not and do express NHA9. Density plots of **d** NHA9 and **e** H3K27ac occupancy at the anchors of annotated CTCF and NHA9 loop anchors. Statistically significant shift detected by the Wilcoxon rank sum test. **f** Density plot of gene expression, log2 (+NHA9/-NHA9), at either CTCF or NHA9 loop anchors. Statistically significant shift detected by the Wilcoxon rank sum test.

To define sets of high-confidence NHA9 and CTCF loops for downstream analysis, we performed CUT&RUN for CTCF and NHA9. NHA9 loops (used throughout this paper) were defined as those that were gained upon NHA9 expression and overlapped NHA9 peaks at both loop anchors, yielding 57 high-confidence “NHA9 loops”, which were detected exclusively in NHA9-expressing cells. Aggregate peak analysis confirmed strong enrichment of Hi-C contacts at these loop positions (**Fig. 1C**). To test whether additional NHA9-mediated contacts might exist below our statistical cutoffs, we examined all potential pairwise interactions between the 1,000 highest occupancy NHA9 peaks, excluding any previously called NHA9 loops, using aggregate peak analysis. This revealed a significant increase in contact frequency in NHA9-expressing cells (Wilcoxon rank sum test p-value < 2.2 × 10^−16^, **Fig. S1A**), suggesting the presence of additional NHA9 loop-like structures exist in these cells that did not meet our threshold for loop detection. For comparison, we defined high-confidence CTCF loops as those that were lost upon acute CTCF degradation and bound by CTCF at both anchors prior to Auxin treatment, yielding a reference set of 7,338 “CTCF loops” for downstream comparisons. As expected, NHA9 loops exhibited a strong enrichment for NHA9 binding compared to CTCF loops (Wilcoxon rank sum test p-val < 2.2 × 10^−16^, **Fig. 1D**).

To explore the functional role of NHA9-mediated loops, we performed CUT&RUN profiling for H3K27ac in HCT116 cells. NHA9 loop anchors exhibited a strong enrichment for H3K27ac signal, with 77% of NHA9 loop anchors overlapping an H3K27ac peak compared to only 15% of CTCF loop anchors. NHA9 loop anchors were also associated with significantly stronger H3K27ac signals compared to CTCF loop anchors (Wilcoxon rank sum test p-val < 2.2 × 10^−16^, **Fig. 1E**). In fact, the 99 highest occupancy H3K27ac sites were all bound by NHA9 (**Fig. S2B-C**). Visual inspection of NHA9 loop anchors revealed extremely broad peaks of H3K27ac often spanning regions of greater than 15 Kb (**Fig. 1A**). H3K27 acetylation increased at NHA9 binding sites upon NHA9 expression, suggesting that NHA9 binding drives H3K27ac deposition rather than NHA9 preferentially binding pre-acetylated loci (Wilcoxon rank sum test p-val < 2.2 × 10^−16^, **Fig. S2A**). Taken together, these findings support the role of NHA9 binding in the formation of broad super-enhancer-like regulatory domains similar to those we previously described in HEK293T cells^12^.

To assess the transcriptional consequences of NHA9-mediated looping, we performed RNA sequencing in both parental and NHA9-expressing HCT116 cells. Genes with promoters located at NHA9 loop anchors were significantly upregulated relative to genes associated with CTCF loop anchors (Wilcoxon rank sum test p-val = 8.3 × 10^−05^, **Fig. 1F**), exhibiting an average increase of 2.26 fold in expression. Upregulated genes at NHA9 loop anchors include known leukemia-associated targets, including *HOXA9, HOXB6*, and *PBX3* (**Supplemental Table 1**). An example of an NHA9 loop that is associated with increased anchor gene expression is shown in **Figure S1B** in which a 80 Kb loop forms connecting a distal H3K27ac peak to the promoter of the gene UBL3, which is upregulated by 5 fold.

Together, these results demonstrate that NHA9 induces a distinct class of chromatin loops in HCT116 cells that connect transcriptionaly active regulatory elements and are associated with gene activation, recapitulating our previous observations in HEK293T cells and establishing that NHA9-mediated chromatin reorganization occurs across multiple cellular contexts.

### NUP98-HOXA9 forms chromatin loops at cell-type-specific locations

Given that NHA9 has only been reported to malignantly transform cells of the hematopoietic lineage^12–17^, understanding whether NHA9 establishes similar chromatin interactions across distinct cellular contexts may provide insight into the lineage specificity of its oncogenic activity. To address this, we compared NHA9-mediated chromatin looping between HCT116 and HEK293T cells to determine the extent to which NHA9 establishes shared versus cell-type-specific chromatin interactions. Because regulatory landscapes and higher-order genome organization differ substantially across cell types, identifying shared and cell-type-specific NHA9 interactions may provide insight into how cellular context influences the genomic interactions established by this oncogenic fusion protein.

To explore the cell-type specificity of NHA9 looping, we compared the NHA9 loops identified in HCT116 cells with previously reported NHA9 loops in HEK293T cells. We reanalyzed our previously published Hi-C and NHA9 ChIP-seq datasets from HEK293T cells expressing either wild-type NHA9 or a phase separation-deficient NHA9 mutant using the same loop-calling and filtering strategy applied in HCT116 cells. This analysis yielded 133 high-confidence NHA9 loops in HEK293T cells. We then took the union of NHA9 loops in HCT116 and HEK293T cells and quantified ‘loop strength’ in each cell type. Loop strength is calculated as the counts in the loop pixel divided by the average counts of local background pixels. Non-looping loci should exhibit scores close to 1, while true loops would have scores substantially higher.

Comparison of NHA9 loops between HCT116 and HEK293T cells revealed that while a subset of loops were shared between cell types, the majority of NHA9 loops are cell-type specific (**Fig. 2A–D**). Only 8% of HEK293T NHA9 loops had loop strengths of greater than 2 in HCT116 cells. Conversely, only 30% HCT116 NHA9 loops had loop strengths of greater than 2 in HEK293T cells. Examples of HEK293T-specific, HCT116-specific, and shared NHA9 loops are shown in **Fig. 2A-C**. Cell type-specific differences in NHA9 looping correlated with cell type-specific differences in gene expression, as genes whose promoters overlapped cell-type–specific NHA9 loops were more highly expressed in their respective cell types (**Fig. 2F**). These data suggest that context-specific NHA9 looping directly impacts differences in gene expression and in turn potentially cellular phenotypes.

**Fig. 2.**
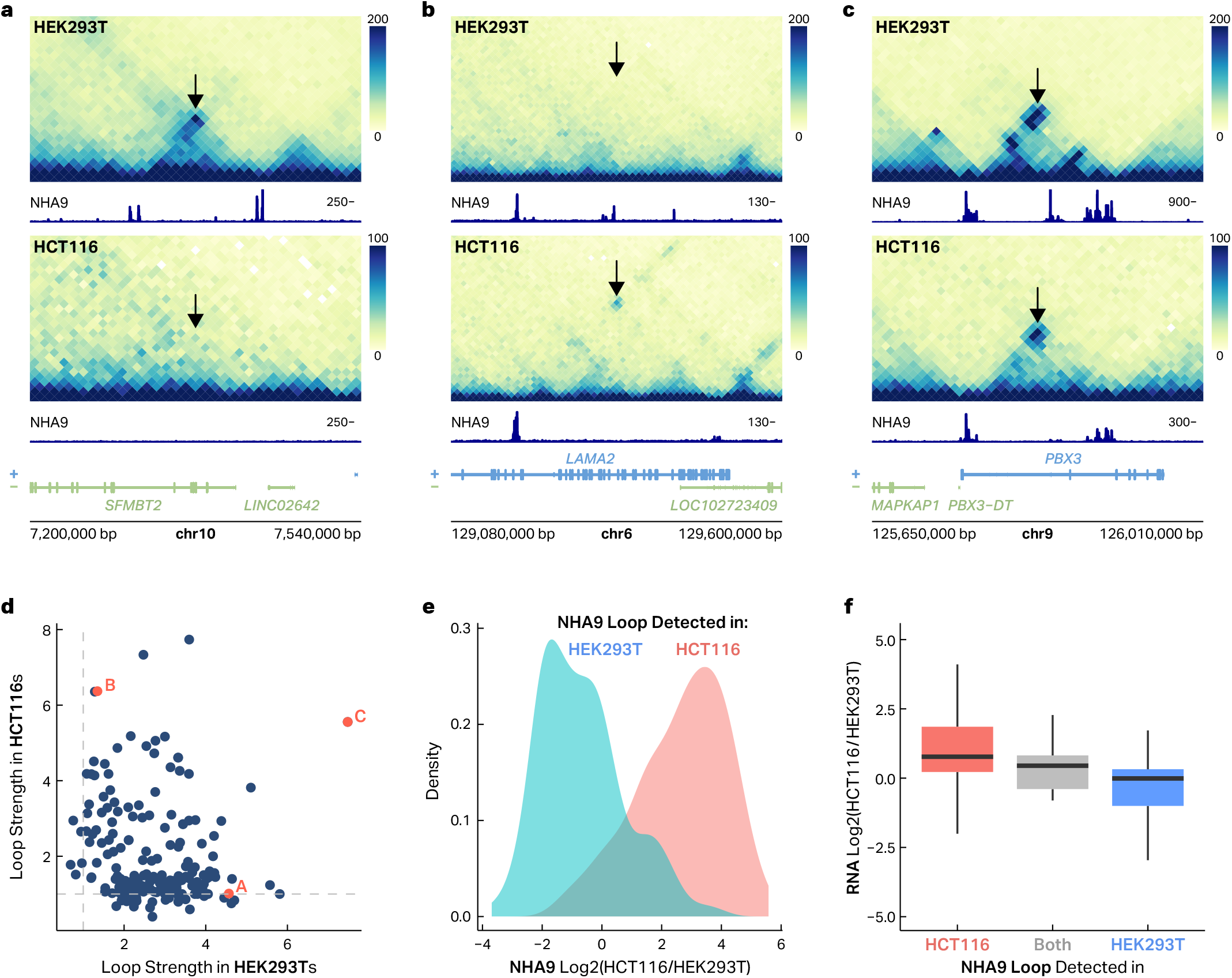
NUP98-HOXA9 forms chromatin loops at cell-type-specific locations. **a**–**c** Hi-C maps of three example regions representing a HEK293T-specific NHA9 loop, a HCT-specific NHA9 loop, and a NHA9 loop present in both cell types. NHA9 signal tracks in their respective cell type are shown below each cell type. **d** Scatter plot of loop strength score for all NHA9 loops in both HCT116 and HEK293T cell lines. **e** Density plot of the log2 fold-change (HCT116/HEK293T) of NHA9 occupancy at NHA9 anchors specific to each cell type. **f** Gene expression log2(HCT116/HEK293T) of genes at cell-type specific NHA9 loops and shared NHA9 loops.

To determine what drives these differences in NHA9 looping we compared these loops to both our NHA9 CUT&RUN data and ChIP-seq data from HCT116 and HEK293T cells. Differences in looping patterns were largely explained by differential NHA9 binding across the two cell lines, as regions engaged in NHA9 looping in one cell type often lacked NHA9 occupancy in the other (**Fig. 2A, 2E**). Only 28% of HEK293T-specific anchors showed NHA9 binding in the HCT116 cell type, and only 16% of HEK293T-specific loops were bound at both anchors in HCT116 cells.

Together, these results indicate that while NHA9 can induce chromatin looping in multiple cellular contexts, the specific interactions that form are largely determined by the underlying, cell-type-specific distribution of NHA9 binding.

### CTCF restricts long-range NUP98-HOXA9 chromatin interactions

Although NHA9 loop anchors are largely not bound by CTCF, it remained unclear whether CTCF is fully dispensable for NHA9 looping or instead constrains the broader NHA9 interaction landscape. To directly test this, we performed *in situ* Hi-C in CTCF-AID HCT116 cells treated with either Auxin to acutely deplete CTCF or with DMSO as a control. As expected, acute CTCF degradation caused widespread loss of chromatin looping. DESeq2 analysis revealed a significant decrease in contact frequency between 8,478 loop anchors including 3,648 of our high-confidence CTCF loops (log2FC < 0, padj < .1, **Fig. 3A-B**). In contrast, NHA9-associated loops were largely retained following CTCF depletion (**Fig. 3A–B**), indicating that NHA9 loop maintenance does not require CTCF. These results highlight that, unlike canonical CTCF-mediated loops, NHA9-driven loops are maintained independently of CTCF. Fluorescence imaging in CTCF-AID cells treated with Auxin or DMSO revealed that NHA9 puncta number, size, and intensity were not significantly altered by CTCF depletion (**Fig. 3C, Fig. S3**), consistent with a model in which NHA9 condensate formation and loop maintenance occur independently of CTCF.

**Fig. 3.**
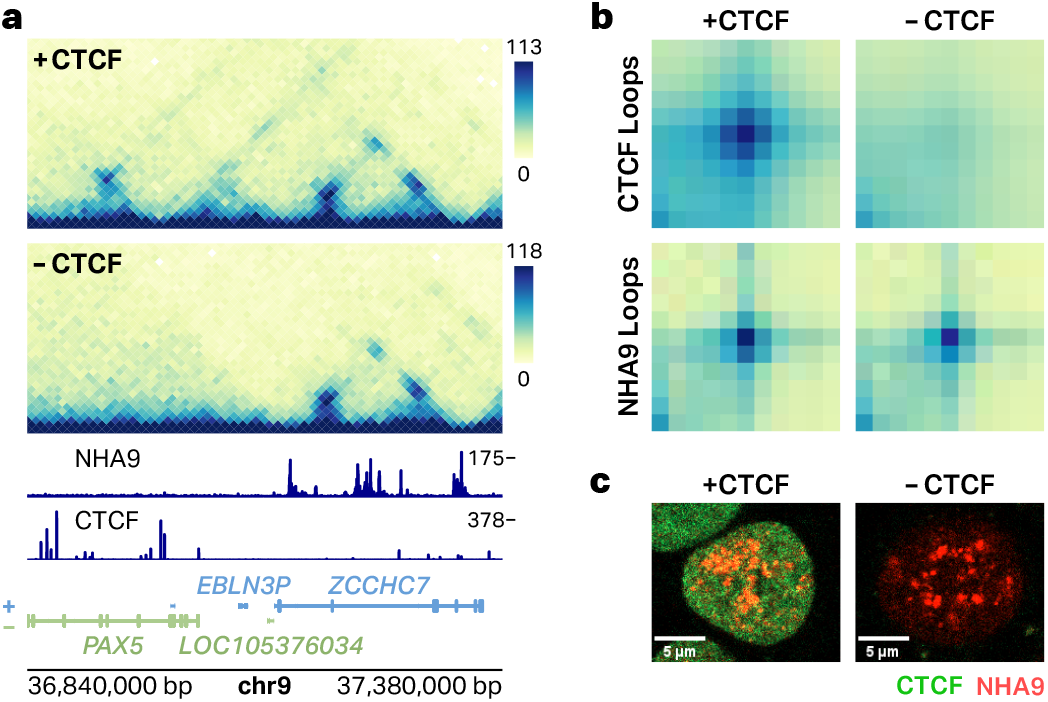
NUP98-HOXA9 loops are independent of CTCF. **a** Representative region showing both CTCF and NHA9 loops in the HCT116-NHA9 cell lines expressing the CTCF degron after treatment with either DMSO or Auxin. Signal tracks for NHA9 and CTCF are shown below in the same region. **b** Aggregate Peak Analysis plots for CTCF and NHA9 loops in HCT116-NHA9 cell lines expressing the CTCF degron after treatment with either DMSO or Auxin. **c** Representative images of the HCT116-NHA9 cells treated with either DMSO or Auxin, with fluorescent imaging of NHA9 and CTCF expressed in the cells.

Upon acute depletion of CTCF, we also observed the emergence of a distinct set of gained chromatin loops **(Fig. 4A)**. These loops are absent in DMSO-treated controls and arise specifically following Auxin-induced degradation of CTCF. Notably, many of these gained loops are bound by NHA9 at one or both anchors and are significantly larger than previously annotated NHA9 loops (Wilcoxon rank sum test p-val < 1.9 × 10^−09^, **Fig. 4B**). These loops are also only detected in HCT116 cells expressing NHA9 and are not observed upon CTCF degradation in the parental HCT116 cell line, indicating that their formation is NHA9-dependent (**Fig. 4C**). We refer to this class of interactions as “restricted NHA9 loops.” Inspection of these 71 restricted NHA9 loops revealed that many connect NHA9-bound regions that were previously segregated into separate CTCF-defined domains. Consistent with this, the NHA9 peaks anchoring these interactions appear spatially constrained under normal conditions and are unable to contact one another in the presence of intact CTCF boundaries. Upon CTCF depletion, these constraints are relieved, enabling the formation of new, long-range NHA9-mediated chromatin interactions **(Fig. 4C)**.

**Fig. 4.**
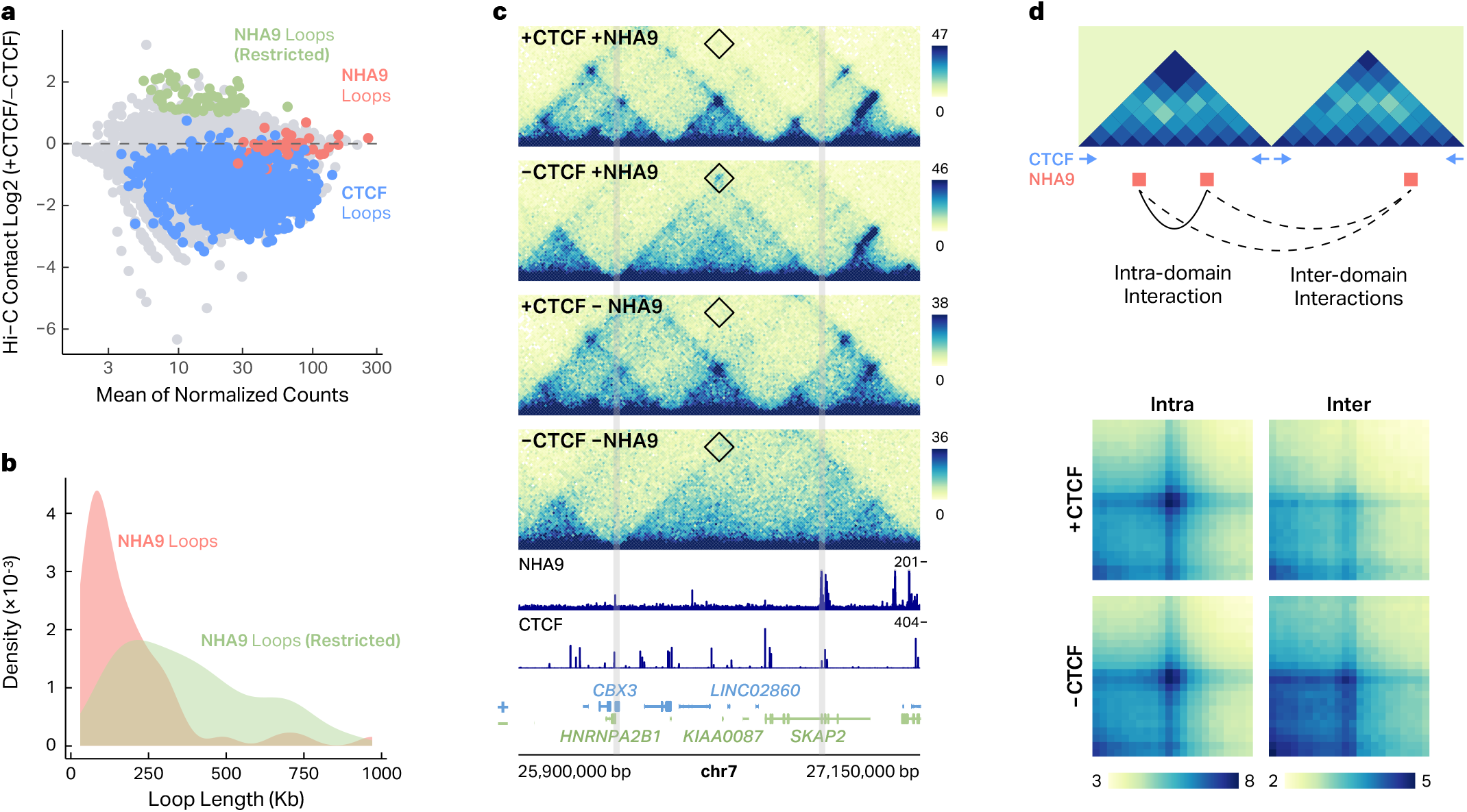
CTCF can restrict the formation of NUP98-HOXA9 chromatin loops. **a** MA plot for all loops after CTCF degradation. CTCF, NHA9, and Restricted loops are annotated in their respective colors. **b** Density plot comparing loop length for NHA9 and Restricted NHA9 Loops. **c** Representative region showing a restricted NHA9 loop with treatment of either DMSO or Auxin in the HCT116-NHA9 cell line expressing the CTCF degron. Signal tracks for both NHA9 and CTCF are shown in the same region. HCT116-Parental cells containing a CTCF degron but not expressing NHA9 are also displayed in the same region with treatment of either DMSO or Auxin. **d** Graphical depictions of what is meant by intra- and interdomain interactions for NHA9 peaks (top). (bottom) APA plots depicting contact frequencies between all intra- and inter-domain pairs of NHA9 binding sites separated by 100-500 Kb reveal an increase in interdomain NHA9 interactions following CTCF degradation.

To directly test whether CTCF domain boundaries limit NHA9-mediated contacts, we quantified pairwise interaction frequencies between NHA9 peaks separated by 150–500 Kb, stratifying interactions based on whether peak pairs reside within the same domain (intra-domain) or across domain (inter-domain) boundaries **(Fig. 4D)**. NHA9 peak pairs located within the same CTCF domain showed no significant change in interaction frequency following CTCF loss. In contrast, NHA9 peaks positioned in adjacent domains exhibited a significant increase (Wilcoxon rank sum test p-val < 3.3 × 10^−13^) in contact frequency upon CTCF degradation **(Fig. 4D)**, with an average fold change of 1.3.

Together, these results demonstrate that while NHA9-mediated chromatin loops form independently of CTCF, their interaction landscape is constrained by CTCF-defined domain architecture. Loss of CTCF relaxes these insulating boundaries, permitting the formation of new NHA9-driven long-range interactions that are otherwise restricted.

### NUP98-HOXA9 loops rely on cohesin in a distance-dependent manner

Because cohesin-mediated loop extrusion underlies most long-range chromatin interactions in mammalian cells, we next asked whether NHA9-mediated loops also depend on cohesin for their formation or maintenance. To test this, we performed *in situ* Hi-C in HCT116-NHA9 cells harboring an AID tag on RAD21 and compared chromatin architecture after treatment with Auxin or DMSO. As expected, RAD21 depletion caused widespread loss of loops, with 10,549 loops exhibiting a significant loss in contact frequency (DESeq2, log2FC < 0 & padj < .1). Consistent with the essential role of cohesin in canonical loop extrusion, 3,803 annotated CTCF loops were significantly lost (DESeq2, log2FC < 0 & padj < .1) upon RAD21 depletion. These results recapitulate results from a number of previous publications confirming the critical role cohesin plays in canonical loop formation and maintenance.

Surprisingly, RAD21 depletion also induced a dramatic weakening of NHA9 interactions (**Fig. 5A–B, SFig. 4A**). 91% (52 of 57) of NHA9 loops exhibited decreased contact frequency in response to RAD21 depletion, with an average log2 fold change of −0.56. 33 of the 57 NHA9 loops met the criteria for a significant decrease in contact frequency while only one met the criteria for a significant increase (DESeq2, padj < .1). Visual inspection (**Fig. 5A**) and aggregate peak analysis (**Fig. 5B**) of NHA9 loops before and after depletion confirm these decreases in contact frequency, revealing that cohesin plays a critical role in NHA9 loop formation and/or maintenance.

**Fig. 5.**
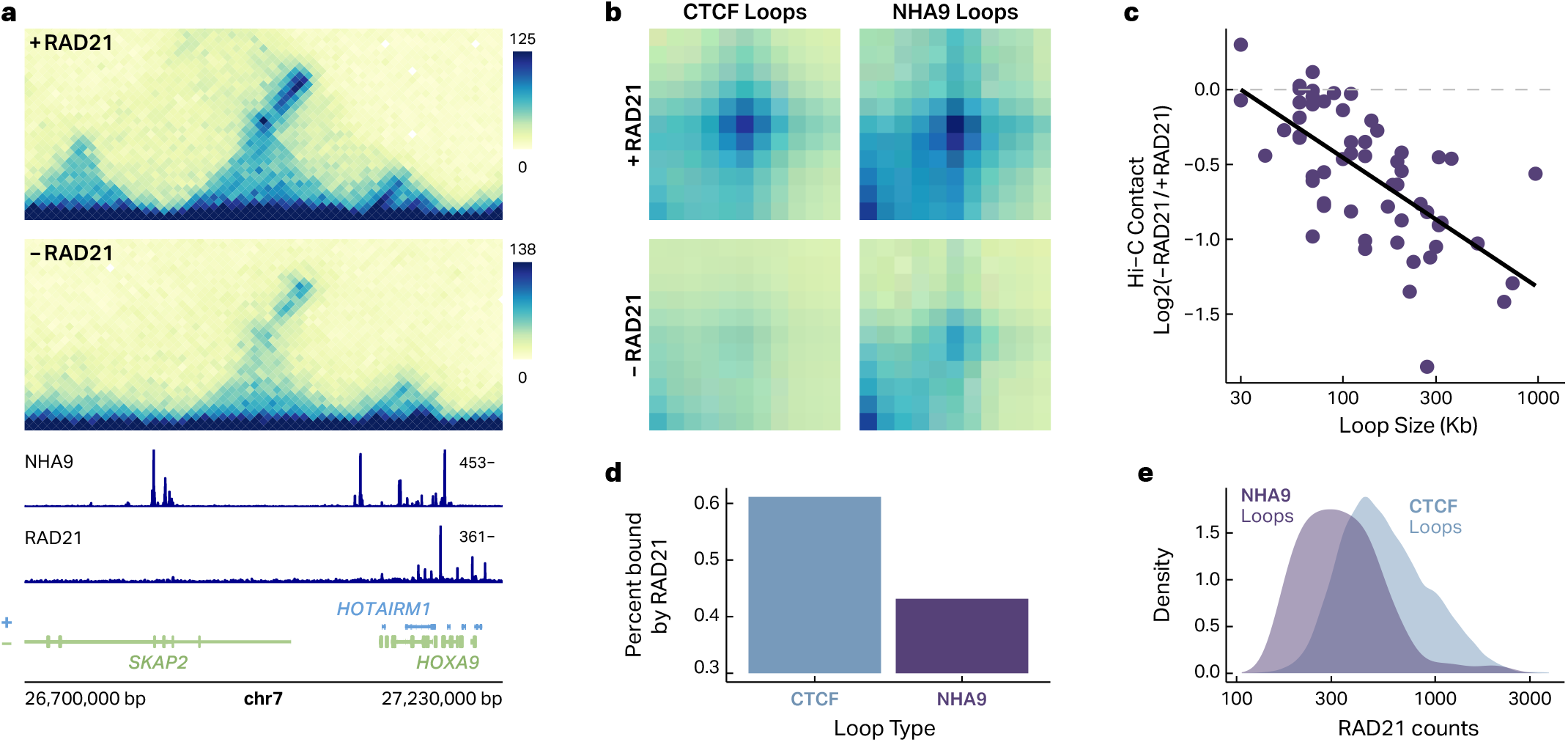
NUP98-HOXA9 loops rely on cohesin in a distance-dependent manner. **a** Representative region showing an NHA9 loop with treatment of either DMSO or Auxin in the HCT116-NHA9 cell line expressing the RAD21 degron. Signal tracks for both NHA9 and RAD21 are shown in the same region. **b** Aggregate peak analysis plots for CTCF and NHA9 loops in the HCT116-NHA9 cell line expressing the RAD21 degron after treatment with either DMSO or Auxin. **c** Scatter plot of log2 change in Hi-C contact frequency of NHA9 loops (Auxin/DMSO) vs loop size in base pairs (bp). **d** Percent of CTCF and NHA9 loops bound by RAD21 at the anchors. **e** Density plots of RAD21 occupancy at the anchors of NHA9 and CTCF loops.

Interestingly, the one NHA9 loop that increased in contact frequency in response to RAD21 depletion was also the shortest NHA9 loop (30 Kb), which prompted us to investigate whether the impact of cohesin on NHA9 loop formation was distance dependent. To address this question, we plotted NHA9 loop size vs the log2 fold-change of contact frequency upon RAD21 depletion **(Fig. 5C)**. RAD21 degradation produced a clear distance-dependent reduction in NHA9 loop strength, with longer loops showing a greater loss of interaction frequency than shorter loops (**Fig. 5C**, r2 = .6). This pattern suggests that cohesin is particularly important for promoting and/or stabilizing long-range NHA9 contacts.

To determine whether cohesin directly anchors NHA9 loops, we examined RAD21 occupancy at NHA9 loop anchors. Only 26% of NHA9 loop anchors overlapped a RAD21 binding site compared to 62% of CTCF loop anchors (**Fig. 5D–E**), suggesting that NHA9 does not function as a direct cohesin barrier in the way that CTCF does. Instead, these findings suggest that cohesin facilitates NHA9 looping indirectly, likely by increasing the probability of contact between distal NHA9-bound loci, interactions that are then maintained by NHA9 phase-separation.

We propose a model in which NUP98–HOXA9 (NHA9) drives a distinct class of phase separation-mediated chromatin loops that are mechanistically separate from canonical CTCF-anchored loops but are shaped by higher-order genome architecture and largely rely on loop extrusion (**Fig. 6**). NHA9 binds specific genomic loci largely independently of CTCF. Instead, CTCF functions primarily as an architectural constraint. While CTCF is dispensable for NHA9 loop formation, CTCF-defined domain boundaries restrict the range of NHA9-mediated interactions by insulating NHA9-bound loci across neighboring domains. When CTCF is removed, these boundaries are relaxed, allowing NHA9 to form new, longer-range “restricted” loops between previously insulated NHA9-bound regions. In contrast, RAD21 is largely required for NHA9 looping, particularly for distal NHA9 binding sites. Cohesin-mediated loop extrusion increases the likelihood of contact between distal NHA9-bound loci, after which NHA9’s phase-separating properties stabilize these contacts into aberrant oncogenic loops. Together, these findings support a hybrid mechanism in which cohesin facilitates chromatin search and proximity, CTCF constrains the permissible interaction space, and NHA9 condensates capture and stabilize selected contacts to create pathogenic chromatin looping networks.

**Fig. 6.**
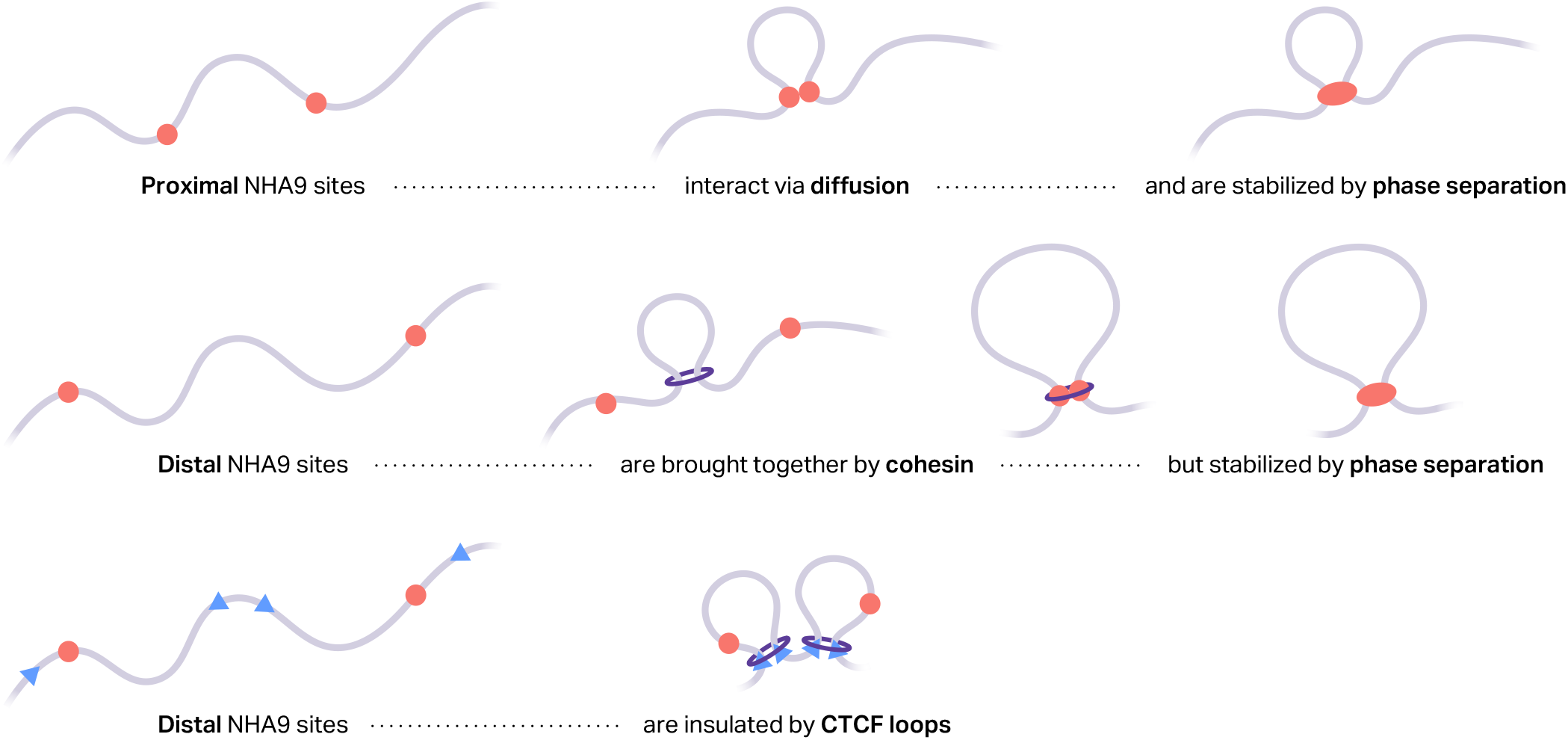
A model of NHA9 loop formation. **top** Interactions between NHA9 binding sites that are close in genomic space occur via diffusion and are stabilized by phase separation. **middle** Initial interactions between NHA9 binding sites that are far apart in genomic space require loop extrusion but are stabilized by phase separation. **bottom** Interactions between NHA9 binding can be ihibitied via insulation by CTCF.

## Discussion

In this study, we investigated how canonical loop extrusion machinery contributes to the formation and organization of NUP98-HOXA9 (NHA9)-mediated chromatin interactions. While our previous work established that NHA9 forms phase separation-dependent chromatin loops associated with enhancer activation and oncogenic transcriptional programs, the relationship between these interactions and canonical looping machinery remained unclear. Here, using engineered human cell lines, we found that NHA9 looping was strongly dependent on cohesin and constrained by CTCF-defined domain architecture, and we propose a model explaining how these factors interact to shape NHA9-mediated chromatin organization. These findings reshape current models of phase separation-driven genome organization by suggesting that condensate-driven chromatin interactions remain fundamentally coupled to canonical architectural machinery. More broadly, this work may have important implications for understanding and therapeutically targeting cancers driven by phase-separating fusion oncoproteins.

By expressing NHA9 in a novel cellular background, we additionally found that NHA9-mediated looping is highly cell-type specific, findings that may help explain the lineage specificity of NHA9-driven leukemogenesis. Previous studies have shown that NHA9 efficiently transforms hematopoietic cells^12–17^, yet its oncogenic activity appears limited outside this lineage. Our data suggest that this specificity may arise, at least in part, from cell-type-specific NHA9 binding and looping. Prior work has implicated factors including Crm1, Menin-MLL complexes, and local chromatin state in regulating NHA9 recruitment and condensate condensation^23–25^; however, the precise epigenetic factors and co-factors required for NHA9 binding to chromatin remain to be established. Nevertheless, these findings support an emerging view that oncogenic fusion proteins function within pre-existing epigenetic frameworks, where lineage-specific chromatin accessibility and cofactor availability shape the downstream oncogenic program beyond simple DNA motif recognition.

While our findings are derived from a single phase-separating fusion protein, they fundamentally reshape current models of phase separation-driven chromatin organization. While prior studies, including ours, have proposed that phase-separating proteins can create chromatin loops by pulling bound loci into nuclear condensates independently of canonical architectural machinery, our data demonstrate that NHA9 looping remains strongly influenced by cohesin-mediated genome organization. In fact, our results suggest that cohesin is still largely responsible for creating the initial contacts, which are then stabilized by phase separation. Because these interactions rely on cohesin extrusion, they are also constrained by CTCF boundaries that impede cohesin processivity. In this framework, phase separation and loop extrusion-driven looping are not opposing mechanisms, but instead function together to shape and stabilize chromatin contact frequencies.

These findings may extend beyond NHA9 biology. Several oncogenic fusion proteins, including EWS-FLI1, PAX3-FOXO1, and BRD4-NUT, have been proposed to reorganize chromatin through condensate formation or aberrant enhancer clustering^19,26–30^. Our findings raise the possibility that many of these fusion-driven chromatin interactions may similarly rely on canonical architectural mechanisms to stabilize or constrain their activity. At the same time, our data suggest that not all non-canonical chromatin interactions will exhibit the same degree of dependence on extrusion. For example, BRD4-NUT-associated megadomains and certain LDB1-mediated interchromosomal interactions likely operate through mechanistically distinct processes, particularly when interactions span chromosome territories that are less compatible with traditional extrusion models^30,31^.

These findings also carry potential therapeutic implications. The dependence of NHA9 looping on cohesin-supported chromatin organization suggests that perturbing architectural pathways could selectively disrupt oncogenic transcriptional hubs even when condensate formation itself is difficult to target pharmacologically. More broadly, therapies aimed at altering chromatin accessibility, enhancer activity, or higher-order genome folding may prove effective in limiting the ability of fusion proteins to establish pathogenic regulatory interactions. As efforts continue to develop therapeutics targeting condensates and transcriptional dependencies in cancer, understanding how phase-separation interfaces with canonical chromatin architecture will likely be critical for identifying the most effective intervention strategies.

Several limitations should be considered when interpreting these findings. First, our analyses rely on engineered HCT116 and HEK293T models that may not fully recapitulate the chromatin environment of primary leukemia cells. Second, although other oncogenic fusion proteins have been proposed to form chromatin interactions through phase separation-related mechanisms, it remains unclear whether they exhibit similar dependencies on cohesin and CTCF. Finally, while our data strongly support a phase-separation mechanism, we do not directly measure condensate biophysics. Our study dissects the consequences of disrupting looping factors on phase separation-driven chromatin looping, but does not directly test the physical and chemical properties of condensates required to create and maintain these structures. Future studies examining NHA9 looping in primary hematopoietic systems, as well as defining the contribution of intrinsically disordered regions and associated cofactors, will be important for further clarifying the molecular basis of these interactions.

Overall, our study provides a mechanistic framework for understanding how phase-separating fusion proteins rewire genome architecture in cancer. By demonstrating that condensate-driven looping remains fundamentally shaped by chromatin context and canonical architectural machinery, this work bridges two major models of genome organization and establishes a foundation for investigating how their interplay contributes to oncogenic gene regulation across diverse cancer types.

## Methods

### Generation of mAID-2 HCT116 Cell Lines expressing NHA9

HCT116 mAID-2 cell lines harboring endogenously tagged CTCF-mAID-2-GFP or RAD21-mAID-2-GFP were obtained from the Kanemaki lab^32^. Cells were cultured at 37 °C, 5% CO_2_, and 5% O_2_ in McCoy’s 5A medium supplemented with 10% fetal bovine serum and penicillin–streptomycin. Stable expression of NHA9 was introduced using the PiggyBac transposon system. Cells at ~50% confluence were co-transfected with the PiggyBac expression plasmid pJA285, encoding NHA9-HA-FLAG-mScarlet under a constitutive promoter and flanked by PiggyBac inverted terminal repeats, together with the PiggyBac transposase plasmid pJA260, using SBI PureFection reagent (cat#LV750A-1). Transfections were performed using a 3:1 expression-to-transposase plasmid ratio. Plasmid preparations were endotoxin-free. Media was replaced 24h post-transfection, and cells were allowed to recover for 2–3 days to permit genomic integration. Stable integrants were selected with puromycin (200 ng/mL) for 7 days and subsequently isolated by fluorescence-activated cell sorting (FACS) on a BD FACSAria™ III cell sorter based on mScarlet expression. Plasmid maps were generated using SnapGene, and full plasmid sequences are provided in the Supplementary Data.

### Cell treatments

HCT116 cells expressing an endogenous CTCF or RAD21 tagged with mAID and GFP that also endogenously express NHA9 tagged with mCherry were grown at 37oC and 5% O2. Cells were treated with 500uM of Auxin (5-phenyl-indole-3-acetic acid, 5-Ph-IAA) alongside control cells treated with an equivalent concentration of DMSO for 6 hours before RNA extraction, CUT&RUN library preparation, or Hi-C library preparation.

### Hi-C library generation

*In-situ* Hi-C libraries were prepared as described by Rao *et al*. Libraries were prepared from 5 million cells cultured to ~80% confluence. Cells were fixed directly in culture medium containing 1% formaldehyde for 10 min at room temperature with gentle mixing, and crosslinking was quenched using glycine to a final concentration of 0.2 M. Cells were washed with cold PBS, scraped from the plates, collected by centrifugation, aliquoted into tubes containing 5 million cells each, flash frozen in liquid nitrogen, and stored at −80°C until processing.

For nuclei isolation, crosslinked pellets were thawed on ice and lysed in Hi-C lysis buffer containing Tris-HCl pH 8.0, NaCl, IGEPAL, and protease inhibitors. Following incubation on ice, nuclei were pelleted, washed, and resuspended in 0.5% SDS prior to incubation at 62°C for chromatin permeabilization. SDS was subsequently quenched with Triton X-100 and water, and chromatin was digested overnight at 37°C using the restriction enzyme MboI in NEBuffer 2 with rotation. After restriction digestion, MboI was heat inactivated and DNA overhangs were filled in using biotin-14-dATP together with dCTP, dGTP, dTTP, and Klenow DNA polymerase to biotinylate fragment ends. Proximity ligation was then carried out in dilute conditions using T4 DNA ligase in the presence of ligation buffer, Triton X-100, and BSA for 4 h at room temperature with slow rotation. Crosslinks were reversed by incubation with proteinase K, SDS, and NaCl, including an overnight incubation at 68°C. DNA was precipitated with ethanol and sodium acetate, washed twice with 70% ethanol, and resuspended in Tris buffer.

Purified DNA was fragmented to an average size of 300–500 bp using a Covaris LE220 focused ultrasonicator. Sonication was performed at a peak incident power of 100, a duty factor of 20, 200 cycles per burst, for 80 seconds. Fragment size selection was performed using sequential AMPure XP bead cleanups to enrich for DNA fragments within the desired size range. Biotinylated ligation products were isolated using Dynabeads MyOne Streptavidin T1 magnetic beads and washed in Tween Wash Buffer at 55°C.

Bead-bound DNA underwent end repair using T4 polynucleotide kinase, T4 DNA polymerase I, and Klenow fragment in ligase buffer supplemented with dNTPs. Following additional washing steps as above, A-tailing was performed using Klenow exo minus and dATP. Illumina indexed adapters were ligated onto repaired DNA fragments using Quick Ligase at room temperature. Adapter-ligated libraries were washed, resuspended in Tris buffer, and amplified directly from beads by PCR using Illumina-compatible primers and Enhanced PCR mix. PCR amplification consisted of an initial denaturation at 95°C followed by 9 amplification cycles with denaturation at 98°C, annealing at 60°C, and extension at 72°C.

Final Hi-C libraries were purified using AMPure XP beads, eluted in Tris buffer, and quantified by Qubit fluorometry prior to sequencing on an Illumina platform. Library quality and fragment size distribution were additionally verified by agarose gel electrophoresis and Bioanalyzer analysis before sequencing. Paired-end sequencing was subsequently performed at Novogene on a NovoSeq 6000 PE150.

### RNA-seq library generation

Following treatments, total RNA was isolated using the RNeasy Mini Kit (Qiagen) according to the manufacturer’s instructions. Briefly, cells were lysed in Buffer RLT supplemented with β-mercaptoethanol to ensure efficient disruption and RNase inactivation. Lysates were homogenized and mixed with ethanol prior to loading onto silica membrane spin columns. Columns were washed sequentially using the provided wash buffers to remove contaminants, and RNA was eluted in RNase-free water.

To minimize genomic DNA contamination, samples were treated with on-column DNase digestion. RNA integrity was evaluated by agarose gel electrophoresis and Bioanalyzer analysis prior to RNA-sequencing.

RNA-sequencing libraries were prepared by Azenta/ Genewiz using standard Illumina-compatible library preparation protocols. BFinal libraries were quality-checked for fragment size distribution and concentration before multiplexed sequencing on an Illumina NovaSeq 6000 platform to generate paired-end reads for downstream transcriptomic analyses.

### ChIP-seq library generation

Chromatin immunoprecipitation (ChIP) was performed as previously described (Bryan et. al 2025 - Dowen lab). All buffers were freshly supplemented with protease inhibitors immediately prior to use. Cell pellets were sequentially lysed to isolate nuclei and prepare chromatin for sonication. Briefly, pellets were resuspended in cold Lysis Buffer 1 (LB1; 50 mM HEPES-KOH pH 7.5, 140 mM NaCl, 1 mM EDTA, 10% glycerol, 0.5% Igepal CA-630, and 0.25% Triton X-100) and rotated at 4°C for 10 min prior to centrifugation at 1,350 × g for 5 min at 4°C. Pellets were then resuspended in Lysis Buffer 2 (LB2; 10 mM Tris-HCl pH 8.0, 200 mM NaCl, 1 mM EDTA, and 0.5 mM EGTA) and rotated for 10 min at room temperature, followed by centrifugation at 1,350 × g for 5 min at 4°C. Pellets were washed once with shearing buffer and finally resuspended in shearing buffer (10 mM Tris-HCl pH 7.6, 1 mM EDTA, and 0.1% SDS) supplemented with protease inhibitors. For spike-in normalization experiments, mouse cells were added at 5% of the total cell number immediately prior to sonication. Chromatin samples were adjusted to a final volume of 1 mL and transferred to Covaris milliTUBEs for shearing. Chromatin was sonicated using a Covaris E220 sonicator with the following settings: duty factor 5%, peak incident power 140 W, 200 cycles per burst, and a total sonication time of 12 min while maintaining sample temperatures between 5–10°C. Following sonication, insoluble debris was removed by centrifugation at 15,000 rpm for 10 min at 4°C. Fragmentation efficiency was assessed by reversing crosslinks on an aliquot of chromatin, purifying DNA using Zymo cleanup columns, and resolving DNA fragments on a 1% agarose gel. Samples displaying chromatin fragments predominantly between 100–700 bp were used for downstream immunoprecipitation.

For immunoprecipitation, Dynabeads were washed in a blocking buffer consisting of 1× PBS supplemented with protease inhibitors and incubated with the RAD21 or IgG antibody for 6–8 h at 4°C with end-over-end rotation. Approximately 5% of each sonicated chromatin sample was reserved as input control prior to immunoprecipitation. Chromatin samples were diluted in post-sonication low-salt buffer (20 mM Tris-HCl pH 8.0, 150 mM NaCl, 2 mM EDTA, 0.1% SDS, and 1% Triton X-100). Additional NaCl and Triton X-100 were added to achieve final concentrations of 300 mM NaCl and 2% Triton X-100 prior to incubation with antibody-conjugated magnetic beads overnight at 4°C with rotation.

Following immunoprecipitation, beads were sequentially washed with post-sonication low-salt buffer, high-salt wash buffer (20 mM Tris-HCl pH 8.0, 500 mM NaCl, 2 mM EDTA, 0.1% SDS, and 1% Triton X-100), LiCl wash buffer (10 mM Tris-HCl pH 8.0, 250 mM LiCl, 1 mM EDTA, and 1% Igepal CA-630), and TE buffer containing 50 mM NaCl (10 mM Tris-HCl pH 8.0, 1 mM EDTA, and 50 mM NaCl). Chromatin was eluted from beads using elution buffer (50 mM Tris-HCl pH 8.0, 10 mM EDTA, and 1% SDS) at 65°C for 1 h with intermittent vortexing to maintain bead suspension. Eluted chromatin and input samples were treated with Proteinase K and incubated overnight at 65°C to reverse crosslinks. DNA was subsequently purified for downstream sequencing.

ChIP-seq libraries were prepared from immunoprecipitated DNA and submitted to the UNC CRISPR Screening Facility for next-generation sequencing. Libraries were sequenced on an Illumina NovaSeq 2000 platform to generate high-depth paired-end sequencing reads for downstream analysis.

### CnR library generation

CUT&RUN experiments were performed using the EpiCypher CUTANA CUT&RUN Kit according to the manufacturer’s protocol. Briefly, cells were harvested and immobilized on activated concanavalin A magnetic beads prior to incubation with primary antibodies directed against the protein or histone modification of interest. Following antibody binding, samples were incubated with pA/G-MNase fusion protein to selectively target chromatin-bound antibody complexes. Targeted chromatin digestion was initiated by calcium activation of MNase, resulting in cleavage and release of DNA fragments associated with antibody-bound chromatin regions. Released DNA fragments were isolated using the purification procedures provided in the kit protocol.

Purified CUT&RUN DNA was quantified and assessed for fragment size distribution prior to library preparation. Sequencing libraries were generated according to the EpiCypher CUTANA workflow recommendations and submitted to the UNC CRISPR Screening Facility for next-generation sequencing. Libraries were sequenced on an Illumina NextSeq 2000 platform to generate paired-end sequencing reads for down-stream genomic analysis.

### Imaging and Quantification

Adherent cells were cultured and prepared for fluorescence imaging on 8-well chamber slides. system. Cells were seeded into chamber wells and allowed to adhere overnight or until approximately 80% confluent. Cells were washed once with 1× PBS and fixed in 4% formaldehyde for 10 min at room temperature.

Formaldehyde was prepared fresh from a 16% stock solution diluted in 1× PBS. Following fixation, wells were washed three times with 1× PBS. Fixed samples could be stored in PBS at 4°C prior to staining.

For immunofluorescence staining, cells were permeabilized in 0.5% Triton X-100 in 1× PBS for 10 min at room temperature and subsequently washed three times with PBS. Samples were blocked in blocking buffer consisting of 2% BSA in 1× PBS for 30 min at room temperature. Primary antibodies were diluted in 0.1% BSA in PBS and incubated with samples either for 3 h at room temperature or overnight at 4°C. Following primary antibody incubation, wells were washed three times with PBS containing 0.05% Tween-20. Fluorescent secondary antibodies diluted in 0.1% BSA in PBS were applied for 1 h at room temperature in the dark. Samples were washed three times with PBS-T followed by a final wash in PBS.

Cells were mounted using fluorescence-compatible mounting medium with DAPI nuclear stain and incubated overnight at room temperature in the dark prior to imaging. Fluorescence imaging was performed using the Zeiss LSM-900.

Cells were imaged on the Zeiss LSM800 confocal microscope using a Plan-Apochromat 63×/1.4 NA oil objective with a 1.7 μm optical section. Images were collected using a 1024 × 1024 field of view at a resolution of 0.10 μm per pixel (101.41 × 101.41 μm FOV), with 16-bits per pixel. The dwell time per pixel was 0.76 μs, with a total frame time of 7.45 s. 488 signal was collected with 0.5% power of a 10 mW laser. 568 signal was collected with 0.5% power for the RAD21 images and 1% power for the CTCF images with 10 mW laser power.

Analysis was done by creating a binary mask for nuclei and entropy per nuclei was calculated using the Matlab entropy function, which provides a numerical value that characterizes the statistical randomness of the inputted array.

### Hi-C Data Processing

In-situ Hi-C datasets were processed using *dietJuicer* (https://github.com/PhanstielLab/dietJuicer), a custom pipeline written in snakemake derived from the Juicer^33^ pipeline. MboI was used as the restriction enzyme, and reads were aligned to the hg38 genome using bwa mem^34^ (v0.7.17). Reads were then filtered to valid pairs with high confidence alignments (MAPQ ≥ 30). Hi-C matrices were constructed for each individual replicate for downstream analysis, as well as a merged map generated for each condition. For visualization, the resulting Hi-C contact matrices were normalized with a matrix balancing algorithm (‘SCALE’) as previously described^6^ to adjust for regional background differences in chromatin accessibility. Hi-C matrices were visualized at specific loci using plotgardener^35^.

### Loop Calling & Differential Loop Analysis

Loops were identified using SIP^36^ (version 1.6.1) with the following settings “-g 2.0 -min 2.0 -max 2.0 -mat 2000 -d 6 -res 5000 -sat 0.01 -t 2000 -nbZero 6 -factor 1 -fdr 0.05 -del true -cpu 1 -isDroso false”. To remove redundancy, loops from all conditions were merged in R with mariner^37^ using the mergePairs function, with merging based on “APScoreAvg”.

A count matrix was prepared using mariner, where unnormalized counts at each loop pixel from each biological replicate were extracted. DESeq2^38^ was used to identify differential loops using the prepared count matrix. Counts from each biological replicate were used with the design “~replicate + condition”. Apeglm was used to calculate log2 fold changes for each loop, comparing the respective conditions. Differential loops were defined as those with an adjusted p-value < 0.1 and an absolute log2(fold-change) > 1, unless otherwise specified.

### Classification of NHA9 and CTCF Loop

Loops were classified as either “NHA9 Loops” or “CTCF Loops” based on the following criteria: NHA9 Loops were required to be (1) gained upon addition of NHA9 (2) both anchors of the loops are bound by NHA9 peaks. CTCF Loops were required to be (1) lost upon degradation of CTCF (2) both anchors of the loops were bound by CTCF peaks. This resulted in 57 NHA9 loops and 7,338 CTCF loops for downstream analysis.

### APA Analysis

Aggregate loop analyses were performed using mariner^37^. Unnormalized Hi-C contact matrices were extracted at 10-kb resolution. Short interactions, loops less than 20kb which would cross the diagonal were excluded, using the removeShortPairs() function. Loop-centric aggregates were generated by centering windows on loop pixels with a 5 pixel buffer. APA plots were generated by the sum of selected regions.

### Loop Strength analysis

Loop strength was quantified as enrichment of the central pixel region relative to a local background, defined as the median of the corner region. Loop enrichment scores were extracted and calculated using the calcLoopEnrichment() in mariner^37^ with the following parameters, FUN = function(fg, bg) median(fg + .01)/median(bg + .01), fg = selectCentrPixel(mhDist = 1, buffer = 10), bg = selectTopLeft(n = 4, buffer = 10) + selectBottomRight(n = 4, buffer =10).

### RNA-seq processing

RNA-seq libraries were processed using the bagPipes snakemake workflow (https://github.com/PhanstielLab/bagPipes/). In brief, read quality was assessed using FastQC^39^ (v0.11.9), adapter trimming was performed with trim_galore^40^ (v0.6.7), and transcript quantification was performed using Salmon^41^ (v1.10.0) against GENCODE hg38 reference indices.

### Differential Gene Analysis

Quantification files were imported into R using tximeta^42^, and counts were summarized to the gene level with tximport. Genes with <10 counts in fewer than 4 samples were excluded. Gene-level count data was analyzed with DESeq2^38^ as previously described. Genes were classified as differentially expressed between conditions if the had an adjusted p-value < 0.1 and an absolute log2(fold-change) > 1, unless otherwise specified.

### ChIP-seq and CUT&RUN processing

ChIP-seq and CUT&RUN libraries were processed using the bagPipes snakemake workflow (https://github.com/PhanstielLab/bagPipes/). In brief, read quality was assessed using FastQC^39^ (v0.11.9), adapter trimming was performed with trim_galore^40^ (v0.6.7), and reads were aligned using bwa mem^34^ (v0.7.17). Samtools^43^ (v1.17) was used to generate a bam file and Picard^44^ (v3.4.0) was used to mark and remove duplicates from the bam file. bigWigs were generated from the filtered bams using deeptools^45^ (v3.5.1) bamCoverage function. Peaks were called on the filtered bam using macs2^46^ (v2.2.9.1) with the following parameters call-peak -f BAM -q 0.01 -g hs --nomodel --shift 0 --extsize 200 --keep-dup all -B --SPMR.

### Differential Peak analysis

A count matrix was generated from the peak calls and bam files using bedtools^47^ (v2.30) multicov. This count matrix was then imported into R and analyzed with DESeq2^38^, as previously described. Peaks were classified as differentially expressed between conditions if the had an adjusted p-value < 0.1 and an absolute log2(fold-change) > 1, unless otherwise specified.

### Signal Enrichment at Loop Anchors

Signal of a protein at loop anchors was determined using the bamCounts() function from the bamsignals library, which determines the number of reads that fall within a genomic region in a bam file.

### Re-analysis of Publicly Available Data

Public Hi-C data from Ahn et al 2021 were downloaded from NCBI Gene Expression Omnibus (GEO) under accession number GSE144643 and processed through the same dietJuicer in-house Snakemake workflow against hg38, using identical alignment (bwa mem^34^ v0.7.17), filtering (MAPQ ≥ 30), SCALE^6^ normalization, SIP^36^ loop-calling parameters, and downstream counting with mariner^37^ (v1.5.0). ChIP-seq datasets for CTCF and H3K27ac in HCT116 cells were downloaded from the ENCODE portal^48^ (https://www.encodeproject.org/) with the following identifiers: ENCSR240PRQ, ENCSR661KMA.

## Supporting information

NHA9 Plasmid Map

## Data Availability

All raw and processed sequencing data generated in this study have been submitted to the NCBI Gene Expression Omnibus (GEO; https://www.ncbi.nlm.nih.gov/geo/). The Hi-C data are available under accession number GSE329508. The RNA-seq data are available under accession number GSE329271. The ChIP-Seq data are available under accession number GSE329273. The CUT&RUN data are available under accession number GSE329275. The code to process and analyze these data is available on GitHub (https://github.com/toobarashid96/MOPS/).

## Author Contributions

T.R. and Z.A.D. conceived the study and designed experiments. T.R. performed analyses, curated data, and wrote the original draft. Z.A.D., D.C.A., G.P., I.Y.Q.-B., S.P., B.K., C.Y. contributed to experiments and data generation. J.D. and W.L. provided methodological expertise, resources, and supervision. D.H.P. provided conceptual guidance, supervision, funding acquisition, and project administration, and served as senior author. All authors reviewed and edited the manuscript.

## Acknowledgements

We thank Erika Deoudes for data visualization, illustration, proofreading, and typesetting. We thank Wendy Salmon and the UNC Hooker Imaging Core. We also thank Brian Golitz and the UNC CRISPR Core for technical assistance.

## Declaration of generative AI and AI-assisted technologies in the manuscript preparation process

During the preparation of this work, the authors used ChatGPT in order to identify weaknesses, correct grammar, and brainstorm titles. After using this tool/ service, the authors reviewed and edited the content as needed and take full responsibility for the content of the published article.

## Supplementary Figures

**Fig. S1.**
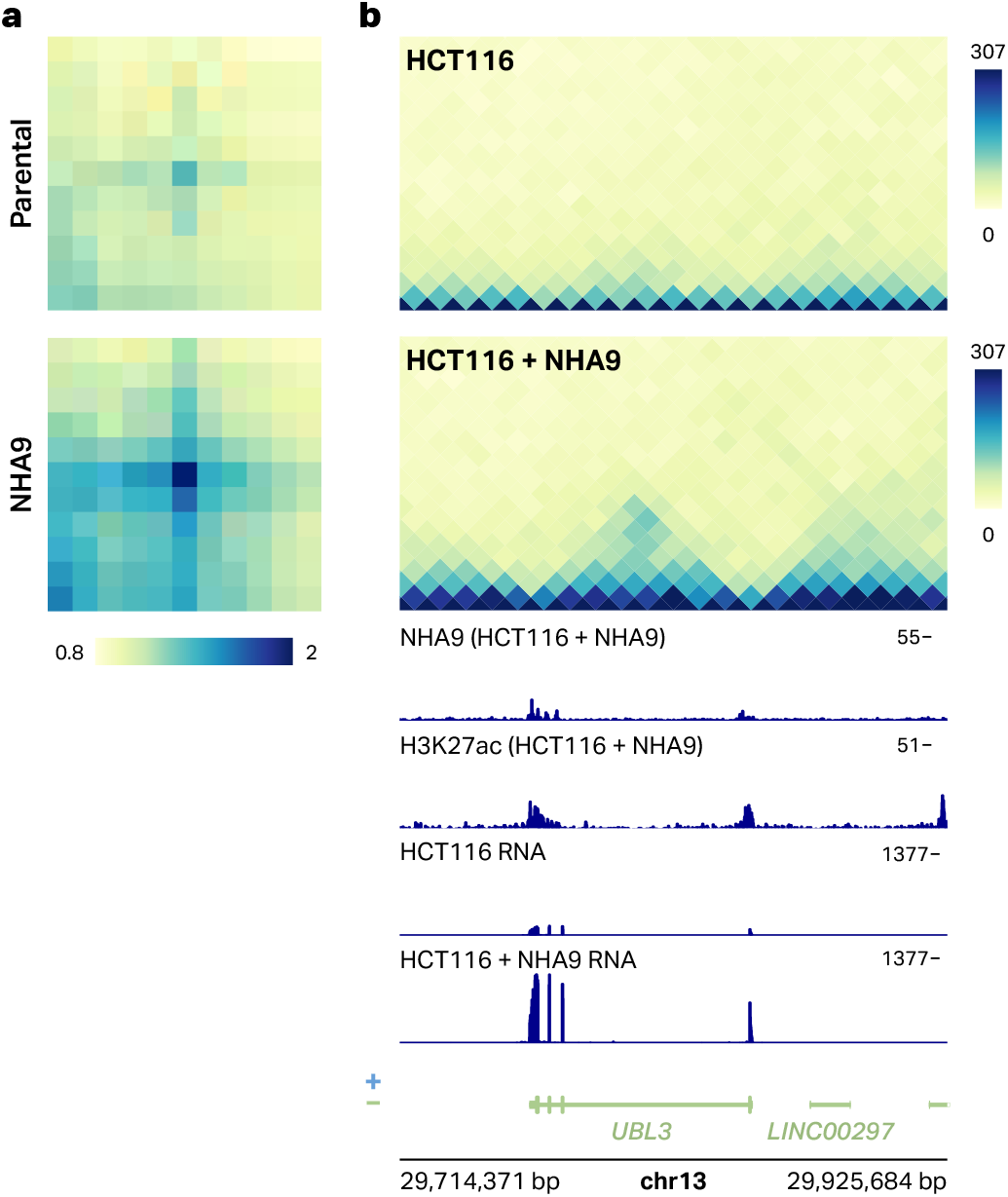
NHA9 binding creates chromatin loops and upregulates loop-associated genes in HCT116 cells. **a**, Aggregate peak analysis plots of all pairwise interactions between NHA9 peaks in HCT116 cells and HCT116 cells expressing NHA9. (B) Example Hi-C region of the UBL3 gene in HCT116 cells +/-NHA9 expression, along with respective signal tracks of NHA9 binding, H3K27ac signal, and reverse-stranded RNA-seq tracks.

**Fig. S2.**
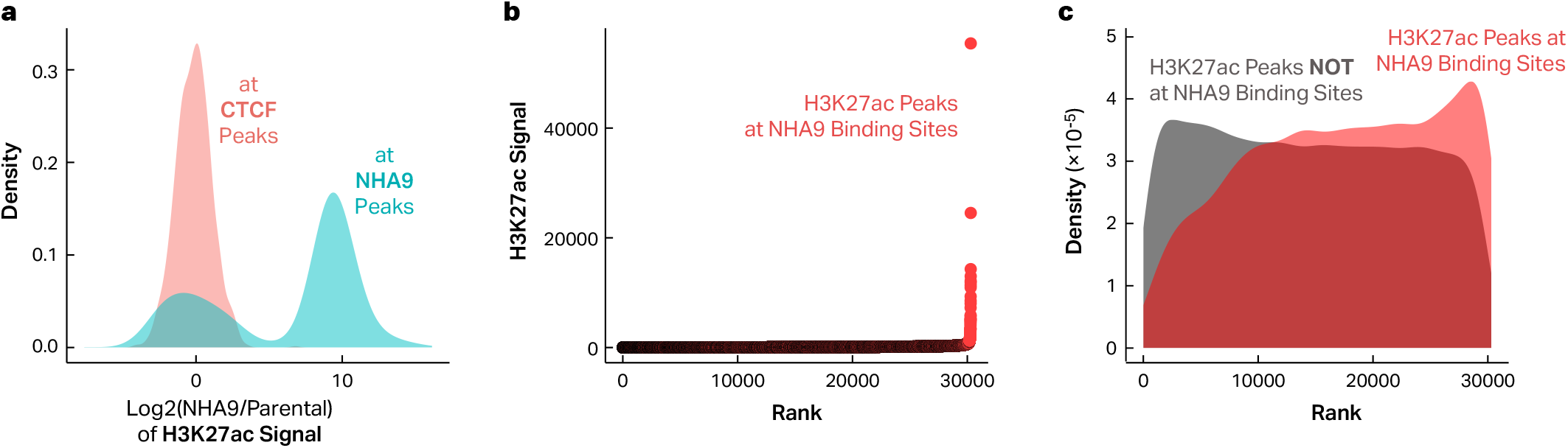
NHA9 expression results in broad, super-enhancer-like H3K27ac binding at NHA9-bound sites. **a**, Density plot comparing the relative amount of H3K27ac signal (NHA9/Parental) bound at CTCF vs NHA9 peaks after the addition of NHA9 expression. (B) Rank-signal plot of H3K27ac peaks, with peaks overlapping an NHA9 peak highlighted in red. (C) Density plot of rank for H3K27ac peaks, comparing those overlapping NHA9 sites to those that are not overlapping NHA9 binding sites.

**Fig. S3.**
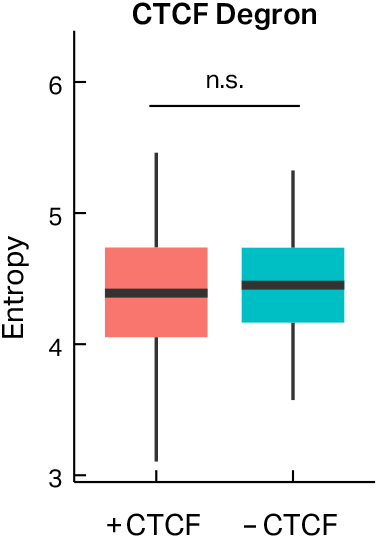
Degradation of CTCF has no impact on NHA9 expression. Boxplots depicting quantified entropy of NHA9 in the nucleus reveal no significant difference between DMSO and Auxin conditions.

**Fig. S4.**
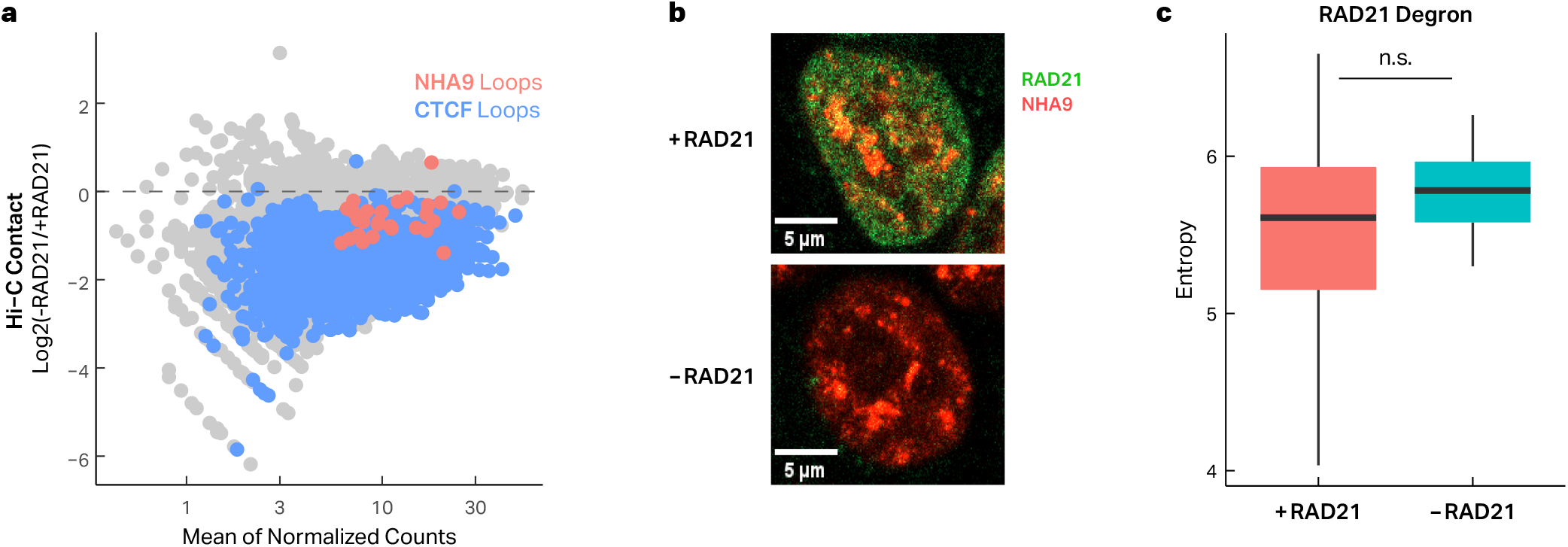
NUP98-HOXA9 loops rely on RAD21 without impacting the expression of NUP98-HOXA9. **a**, MA plot for all loops after RAD21 degradation, comparing the log2FC (Auxin/DMSO). CTCF and NHA9 loops are annotated in their respective colors. (B) Representative images of the HCT116-NHA9 cells treated with either DMSO or Auxin, with fluorescent imaging of NHA9 and RAD21 expressed in the cells. (D) Entropy quantification of NHA9 in the nucleus represents relative NHA9 expression in the DMSO and Auxin conditions.

