## Supplementary figures and images for "Aberrant chromatin looping by NUP98-HOXA9 is constrained by CTCF and facilitated by cohesin"

### NHA9 Plasmid Map

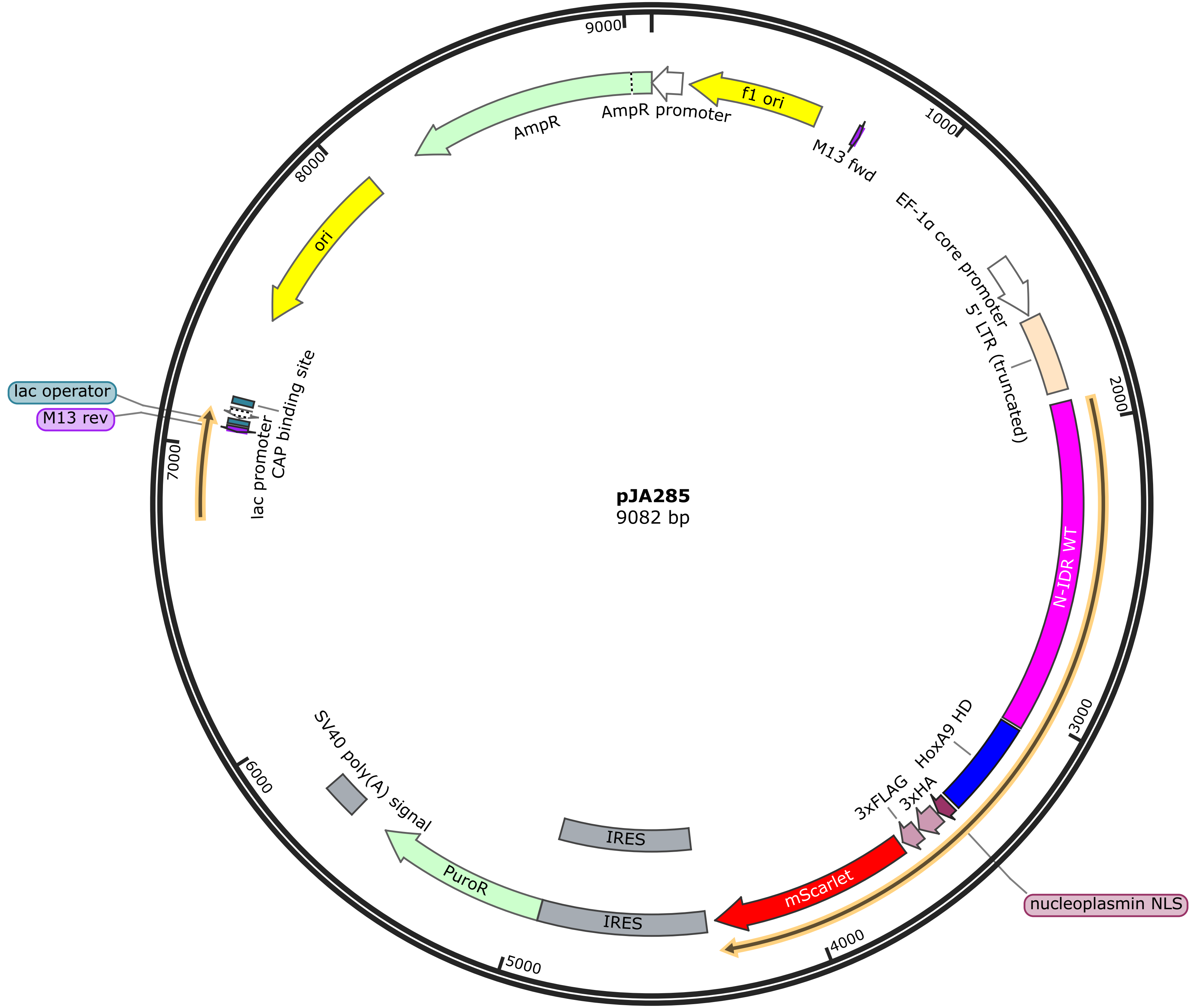
